# Host independent deletion hotspots in the SARS-CoV-2 genome

**DOI:** 10.1101/2022.10.16.512395

**Authors:** Mohammad Khalid

## Abstract

SARS-CoV-2 infects a wide range of hosts in varying degrees. The RNA genome of SARS-CoV-2 makes it prone to mutations. Advantageous mutations help the virus to evolve and the virus maintains such mutations across species. Here in this study, all non-human hosts-derived SARS-CoV-2 genomic sequences from the GISAID database were analyzed, and identified several deletion hotspots, which are maintained by the virus, across various host species, indicating their important role in the virus evolution. Several of these deletion hotspots are also found in human-derived SARS-CoV-2 genomic sequences. These deletion hotspots have the potential to affect the pathogenicity and virulence of the virus and have a role in molecular and serological diagnostics. Potentially, they can lead to immune escape, resulting in vaccine failure and drug-resistant variants.

## Introduction

In December 2019, an outbreak of SARS-CoV-2 led to COVID-19 pandemic, which caused unprecedented damage on healthcare and socio-economic fronts. The virus is transmitted to human by zoonotic spillover (1) may be from bat (2) or pangolin (3). The virus uses Spike protein to interact ACE-2 receptor on host cell surface to infect the cell (4-6). Molecular and serological techniques have been employed to detect the virus using nasopharyngeal samples.

The SARS-CoV-2 has a positive-sense single-stranded RNA (+ssRNA) genome of 29-30 Kb size (7). The RNA genome makes the virus more prone to mutations due to lack of proofreading function of RNA-dependent RNA polymerase (RdRp) (8). However, viruses from coronavirus family are known to have a proofreading mechanism (9, 10) expecting to have low mutation rate, though, thousands of mutant variants of the virus have been emerged over the span of two years, infecting people across the globe (11). Some of these mutations are able to generate specific genetic lineages that have outcompeted the original virus (12). These lineages are characterized by higher transmission rate or virulence or reduced effectiveness of counter measures and classified by WHO as variants of concern (VOCs) (13).

Mutational instances occurring at same genomic locations in variety of hosts are mutational hotspots, potentially, they make viral evolution incredibly repeatable across the hosts. SARS-CoV-2 could evolve along these hotspots, driving its evolution to make the virus more efficient in transmissibility and virulence (14, 15) Furthermore, such hotspots might affect primer-binding and probe-binding to target sites, and neutralizing antibodies binding, which can reduce the sensitivity of molecular and serological diagnostic tests (16, 17). Moreover, these mutations can cause vaccine failure and emergence of antiviral drug resistant viral variants.

Zoonotic spillover of the SARS-CoV-2 has motivated this study to analyze the SARS-CoV-2 genomic sequences, derived from non-human hosts to investigate common mutations in the viral genome and to identify deletion hotspots across the viral hosts.

## Methods

### SARS-CoV-2 genome sequences derived from non-human hosts

In this study, all 2,162 available full-length SARS-CoV-2 genome sequences derived from non-human hosts obtained from the GISAID database (11) as of 27^th^ August 2022 were analyzed. These SARS-CoV-2 genomic sequences were derived from all non-human hosts across the globe viz; mink, deer, cat, dog, lion, tiger, hamster, mouse, bat, gorilla, pangolin snow leopard, otter and monkey.

### Analysis of deletion mutations in non-human hosts derived SARS-CoV-2 genome

The FASTA format file of the SARS-CoV-2 genome sequences were downloaded from GISAID database and analyzed using online tools Nextclade of Nextstrain (18-21). Nextclade uses SARS-CoV-2 Wuhan-Hu-1/2019 (Accession # MN908947) isolate as a reference genome. The Algorithm was Sequence alignment, Translation, Mutation calling, Detection of PCR primer changes, Phylogenetic placement, Clade assignment and Quality Control. The data were compiled and presented using Microsoft Excel 2019.

Every deletion in nucleotide sequences from reference strain, considered as a mutation if the deletion happened more than one instance in the genome.

### Phylogenetic analysis of non-human derived SARS-CoV-2

The phylogenetic tree was generated by NextStrain Auspice (18). The phylogenetic tree was visualized from NexStrain inferred evolutionary relationships between different viral genomes derived from non-human hosts, based on their genomic sequences.

## Results

### Hosts and their number of genomic sequences

The details of the SARS-CoV-2 genome sequences derived from all non-human hosts and submitted to GISAID database (11). The viral sequences were available as on 27 August 2022. Mink (*Neovison vison*) seems to be susceptible to the infection and has second highest number of genomic sequences derived from it, as shown in the Table 1.

**Table 1:** The SARS-CoV-2 infection in various hosts.

| S. No. | Name | Sequences |
| --- | --- | --- |
| 1 | Human (*Homo sapiens*) | 12,590,689 |
| 2 | Mink (*Neovison vison*) | 1,346 |
| 3 | Deer (*Odocoileus virginianus*) | 332 |
| 4 | Cat (*Felis catus*) | 121 |
| 5 | Dog (*Canis lupus familiaris*) | 84 |
| 6 | Lion (*Panthera leo*) | 75 |
| 7 | Mink2 (*Neogale vison*) | 45 |
| 8 | Tiger (*Panthera tigris*) | 42 |
| 9 | Hamster (*Mesocricetus auratus*) | 30 |
| 10 | Mouse (*Mus musculus*) | 23 |
| 11 | Bat (*Rhinolophus*) | 15 |
| 12 | Gorilla (*Gorilla gorilla*) | 15 |
| 13 | Pangolin (*Manis javanica*) | 13 |
| 14 | Snow Leopard (*Panthera uncia*) | 9 |
| 15 | Otter (*Aonyx cinereus*) | 8 |
| 16 | Monkey (*Chlorocebus sabaeus*) | 4 |

List of SARS-CoV-2 genome sequences on GISAID database from various hosts, available as on 27/08/2022.

### Phylogenetic tree of non-human derived SARS-CoV-2

Phylogenetic tree generated with NextStrain analysis tool using non-human hosts derived genomic sequences from the GISAID database, as shown in Figure 1. The figure also revealed clad information of these genomic sequences.

**Figure 1:**
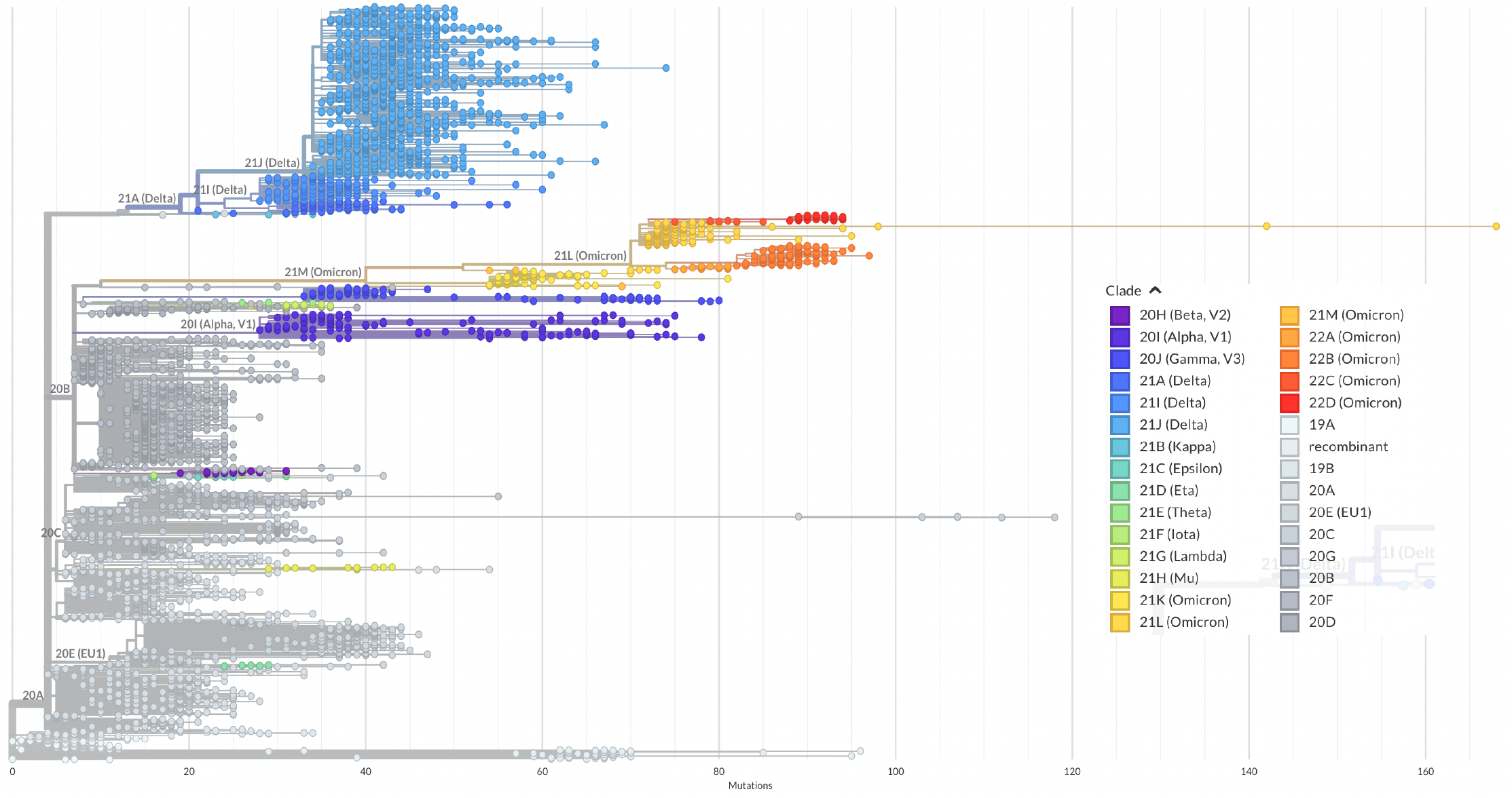
Phylogenetic tree of the SARS-CoV-2 genomes, derived from various hosts. Phylogenetic tree of circulating of SARS-CoV-2 genomic sequences derived from non-human hosts. The phylogenetic tree was generated using the NextStrain Auspice online tools.

Bat and pangolin derived genomic sequences excluded from the phylogenetic analysis because they behaved very differently; placing themselves far away from the rest of the genomic sequences and that hid details of the remaining samples.

### Deletion mutations in non-human derived SARS-CoV-2

During the analysis of 2,162 full-length SARS-CoV-2 genome sequences, in this study 187 deletion mutations have been found across all the non-human hosts (Supplementary Table 1). Out of 2162 samples, 4 samples were derived from monkey (Table 1) but none of these 4 samples have deletion when compared to reference genome (Accession # MN908947). Mouse derived genomic sequences contributed 23 samples and contain many deletions but they were single instances so discarded because of the study criteria, only one deletion mutation (26260-26283) was found in mouse derived samples and that was unique to mouse only (Supplementary Table 1). Bat and pangolin derived genomic sequences had many deletion mutations (73 and 35 respectively) and most of them are unique to either bat or pangolin only (Supplementary Table 1).

### Identification of deletion hotspots across the hosts

Supplementary Table 1 revealed many deletion mutations are common across the hosts in various ORFs. These 10 common deletion locations are the hotspot of deletion mutations viz; 516-518, 6506-6508, 6513-6515 and 11288-11296 are laying in ORF1a, while 21765-21770, 21992-21994, and 22029-22034 are placed in Spike gene and 28248-28253, are laying in ORF-8, 28271 is in between ORF-8 and N, and 28362-28370 in ORF-N. It is very less likely that these hosts were infected by the common variants of the virus, because the genomic sequences were collected from these hosts at various different time point as well as from different geographical locations. These 10 deletion hotspots were analyzed for the occurrence in every host, Figure 2. From these 10 deletion hotspots, 6 are found in majority of the hosts in high percentage, these deletion hotspots are 11288-11296, 21765-21770, 21992-21994, 22029-22034, 28248-28253, and 28271.

**Figure 2:**
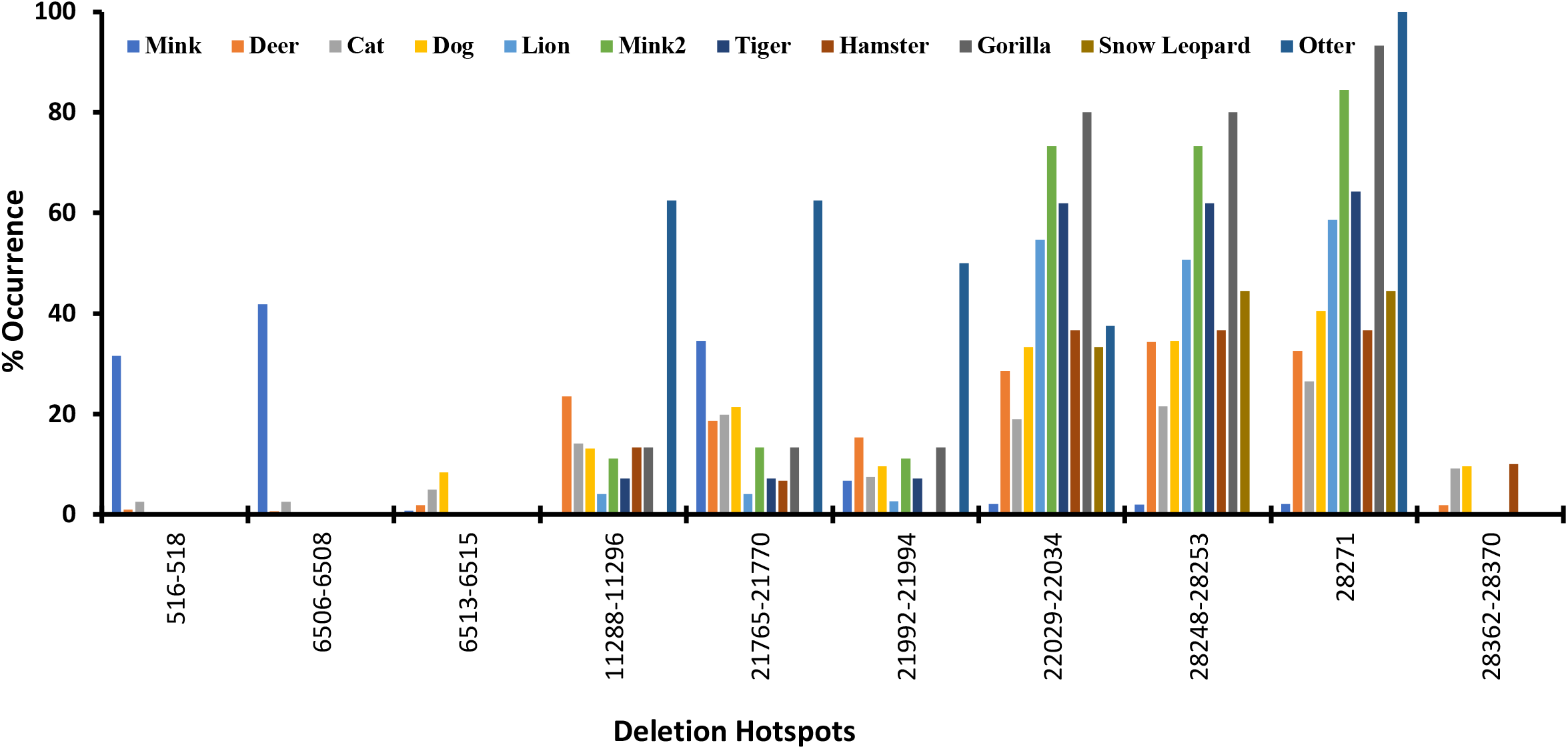
Deletions hotspots and their occurrence in various host. Identification of deletion hotspots in SARS-CoV-2 genome derived from non-human hosts. The analysis based on all SARS-CoV-2 genome sequences derived from non-human hosts, available on GISAID database as on 27/08/2022.

### List of identified deletion hotspots and position in ORFs: Table 2

The 10 deletion hotspots; 516-518, 6506-6508, 6513-6515 and 11288-11296 are laying in ORF1a, while 21765-21770, 21992-21994, and 22029-22034 are placed in Spike gene and 28248-28253, are laying in ORF-8 and 28271, and 28362-28370 in ORF-N, are shown in Figure 3 (not on scale) and amino acid truncation are also listed, Table 3.

**Table 2:** Deletion hotspots and their amino acid truncation.

| Mutation No. | Deletion Position | ORFs | AA Deletion |
| --- | --- | --- | --- |
| 11 | 516-518 | 1a | 1a: NSP1: M85Δ |
| 33 | 6506-6508 |  | 1a: T2081Δ |
| 35 | 6513-6515 |  | 1a: S2083Δ |
| 40 | 11288-11296 |  | 1a: NSP6: SGF 3675–3677Δ |
| 58 | 21765-21770 | S | S: H69/V70 Δ |
| 73 | 21992-21994 |  | S: Y144Δ |
| 78 | 22029-22034 |  | S: EF156-157Δ |
| 166 | 28248-28253 | 8 | ORF8: DF119-120Δ |
| 171 | 28271 |  | M1Δ |
| 174 | 28362-28370 | N | N: ENAΔ |

List of deletion hotspots in the SARS-CoV-2 genome. The SARS-CoV-2 genome sequences derived from non-human hosts, available on GISAID database as on 27/08/2022.

**Figure 3:**
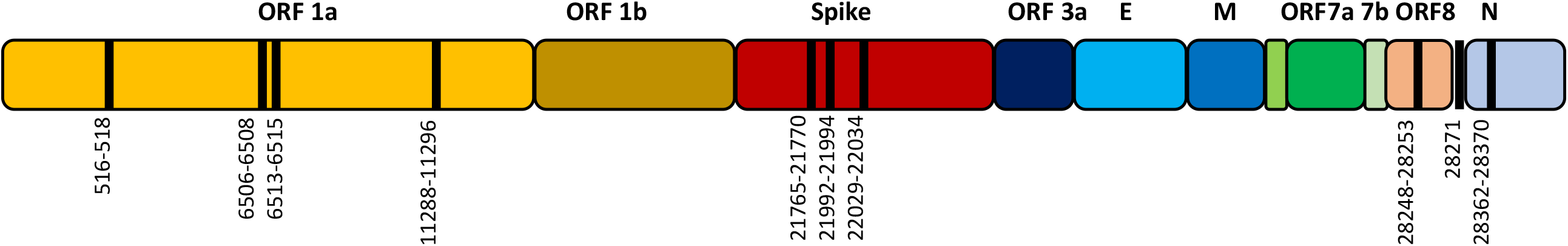
Deletion hotspots. Schematic diagram of SARS-CoV-2 genome with various ORFs. The black line representing deletion hotspots in the SARS-CoV-2 genome (not on scale).

## Discussions

The spike protein of SARS-CoV-2 interacts with the ACE-2 receptor in the host’s cell. The interaction of the spike protein with the ACE-2 makes a cell prone to the virus infection (4-6). Spike protein interacts with ACE-2 from various species, allowing the virus to cross species barriers, as shown in Table 1, the SARS-CoV-2 is capable of infecting a wide range of mammalian hosts. Mink (*Neovison vison*) derived virus genomic sequence is second highest, only to that of humans, and the virus has been found to cause viral pneumonia in the mink (22, 23). The SARS-CoV-2 isolates derived from mink and humans also found to have several common mutations (24).

The SARS-CoV-2 outbreak is a zoonotic spillover suspected to be from bat or pangolin (1-3). All 2162, non-human derived SARS-COV-2 genomic sequences were analyzed for genomic similarities with reference to Wuhan-Hu-1/2019 (Accession # MN908947) isolate. The bat and pangolin derived SARS-COV-2 genomic sequences were not closely related, as a result, phylogenetic tree overlooked the information about the remaining genomic sequences (data not shown), and so the bat and pangolin derived SARS-COV-2 genomic sequences were excluded from the phylogenetic analysis (Figure 1).

The non-human derived genomic sequences belong to variety of clads but majority of these samples were from clad 20A, B, C, E and 21J (Delta), suggesting variety of clades of SARS-CoV-2 was transmitted to variety of non-human hosts, Figure 1.

The SARS-CoV-2 genome is very plastic to accommodate variety of mutations; from the 2162 genome sequence analysis revealed 187 deletion mutations (Supplementary Table 1). Several of them are unique to specific hosts; the SARS-CoV-2 genomic sequences derived from bats contained the highest number of deletion mutations (73), most of them were unique to bat derived samples only, suggesting non-advantageous deletions. While many other mutations were found across the mammalian species and considered deletion hotspots in this study, 10 of them are worth mentioning viz; 516-518 is a deletion in NSP1 at position M85Δ, found in mink, deer and cat derived SARS-CoV-2 genomic sequences, which was also reported to be in human-derived samples (25, 27). Deletion 6506-6508 is translated to ORF1a T2081Δ, found in mink, deer and cat derived samples. Another ORF1a deletion, 6513-6515 is translated to S2083Δ found in this study in min, deer, cat and dog derived SARS-CoV-2, which was also reported in human-derived SARS-CoV-2 genomic sequences in Vanmechelen et. al., study (26). The deletion 11288-11296 in ORF1a: NSP6: SGF 3675–3677Δ found in this study, was also reported in human derived genomic sequences (27). While 21765-21770 deletion is in S: H69/V70Δ which was reported in human derived samples, causing increased viral transmission and infectivity and the virus variants was known as Omicron was also found in human derived samples and became one of the dominant variants across the globe (26, 27, 28). The 21992-21994 and 22029-22034 deletions reported in this study in majority of hosts derived genomic sequences, were also reported in human-derived SARS-CoV-2 genome sequences in non-RBD S1 region and known to have role in viral transmission and infectivity (27, 28). The deletion 28248-28253 translated as DF119-120Δ in ORF8 reported here, which was also found in human-derived SARS-CoV-2 genome sequences (27). The 28271 deletion is nucleotide deletion in non-coding region between ORF-8 and N reported in this study in most of the non-human host derived SARS-CoV-2, was also found in human derived samples (26, 27), and 28362-28370 is deletion in N: ENA27-29Δ was also reported to be found in human-derived SARS-CoV-2 genome sequences (26).

This study reports 10 deletion hotspots in SARS-CoV-2 genome sequences, of those 7 hotspots were found in most of the hosts (Supplementary Table 1 and Figure 2) and also reported in human derived SARS-CoV-2 genome sequences in various studies. Many of these deletion hotspots have functional role in the virus pathogenicity and virulence.

## Conclusions

The SARS-CoV-2 has a modest rate of mutation, these mutations help the virus to evolve to counter hosts defenses. The virus conserves many of such deletion mutations at same genomic locations, irrespective of hosts. Such hotspots have potential to decrease the efficiency of molecular and serological diagnostic tests as well as failure of vaccines and antivirus drugs.

## Acknowledgment

I would like to extend my thanks to the people at NCBI database and GISAID for making the SARS-CoV-2 genome sequences available.

## Abbreviations

GISAID: Global initiative on sharing all influenza data
NSP: Non-structural protein
ORF-8: Open Reading Frame-8
UTRs: Untranslated regions

